# What’s Up: an assessment of Causal Inference in the Perception of Verticality

**DOI:** 10.1101/189985

**Authors:** K.N. de Winkel, M. Katliar, D. Diers, H.H. Büelthoff

## Abstract

The perceptual upright is thought to be constructed by the central nervous system (CNS) as a vector sum; by combining estimates on the upright provided by the visual system and the body’s inertial sensors with prior knowledge that the upright is usually above the head. Results from a number of recent studies furthermore show that the weighting of the respective sensory signals is proportional to their reliability, consistent with a Bayesian interpretation of the idea of a vector sum (Forced Fusion, FF). However, findings from a study conducted in partial gravity suggest that the CNS may rely on a single sensory system (Cue Capture, CC), or choose to process sensory signals differently based on inferred signal causality (Causal Inference, CI). We developed a novel Alternative-Reality system to manipulate visual and physical tilt independently, and tasked participants (n=28) to indicate the perceived upright for various (in-)congruent combinations of visual-inertial stimuli. Overall, the data appear best explained by the FF model. However, an evaluation of individual data reveals considerable variability, favoring different models in about equal proportions of participants (FF, n=12; CI, n=7, CC, n=9). Given the observed variability, we conclude that the notion of a vector sum does not provide a comprehensive explanation of the perception of the upright.

## 1 Introduction

Whenever we specify objects’ relative locations using terms as ‘above’ or ‘below’, or when we move about throughout the world while trying not to topple over, we make use of the fact that we have a perception of upright. Multiple sensory systems throughout the body provide the nervous system with information that can potentially be used to construct a subjective vertical: visually, we are able to determine our orientation from polarity information in the optic array[1]; our vestibular system is stimulated by accelerations, and therefore provides us with information on the direction and magnitude of the gravitational vector[2]; we receive information about our orientation relative to gravity from pressure cues on the body[3, 4] and the distribution of fluids in the body[5, 6]; and there is evidence for specialized graviceptors located in the trunk[7]. Mittelstaedt [8] proposed that the Central Nervous System (CNS) constructs perceptions of verticality by combining the sensory information from the visual system and the body’s collective inertial sensors with the prior knowledge that ‘up’ is usually aligned with the long-body axis (the idiotropic vector), and that the process could be described as a vector sum where the length of the vectors represents the relative influence of each component. In subsequent work, this concept has been interpreted as a reflection of statistically optimal behavior by the Central Nervous System (CNS): according to Bayes’ rule, if sensory estimates of the upright are normally distributed random variables and the prior is either normally distributed or uninformative, the estimate that is most likely the true upright can be calculated as a weighted average of the sensory estimates, where the weights are proportional to the inverse of estimates’ variances [9, 10].

Several studies indeed report that people’s perception of the upright reflects such Bayesian integration [11, 12, 13, 14]. Different measures were used among studies: participants were instructed to either indicate the Subjective Visual Vertical (SVV) by aligning an object in the visual display with the perceived upright[11, 12, 13, 14]; the perceptual upright was inferred from participants’ interpretations of the ambiguous symbol ‘p’, which is defined by its orientation relative to the perceived upright (equivalently, ‘d’)[11, 15]; or participants indicated Subjective Body Tilt[13](SBT). For the former measures, it was found that the calculated ratios of the weights attributed to the constituent cues coincided well with calculations of sensory variance, thereby providing supporting evidence for the Bayesian interpretation of the vector-sum model; for the latter measure, this evidence was obtained by comparisons of model fit indices.

However, some findings are inconsistent with such modeling. First, the weightings reported differ between experiments. From the perspective of a Bayesian vector sum model, this suggests that the sensory variances differ between tasks and experiments. Even though it is not unlikely that specific conditions of an experiment affect the sensory estimates, it is not clear why the variance of sensory estimates of the upright would vary depending on the task. Second, [16] performed a study where participants were asked to report the SVV during exposure to different levels of gravity during parabolic flight. Here, it was found that participants discarded the visual cue entirely, and relied on either inertial or idiotropic information in a dichotomous fashion, where the probability of landing on the inertial cue was proportional to the strength of the gravitational pull. Similarly, [17] found hardly any distortion at all in an SBT task, which was interpreted as an indication that the idiotropic vector affects the SVV, but not SBT.

The observed differences in the role specific cues fulfill in different tasks and the aforementioned variability in sensory weightings imply that Bayesian models based on the notion of a vector sum cannot offer a comprehensive explanation of the perception of the upright. It is possible that the role attributed to different cues reflects the inferred causality of the signals, in relation to the particular task. A class of models that can account for different behaviors are Causal Inference (CI) models. According to these models, assessments of causality are included in the formation of perceptions. Put simply, these models state that the CNS constructs intermediate estimates of environmental properties consistent with different interpretations of their causes (i.e., a common cause or separate causes) in tandem, and combines these into final estimates, taking into account the probability of the respective causal structures. We hypothesized that CI models provide a better explanation of the perceptual upright than the vector-sum approach. In an experiment, we independently manipulated participants’ physical and visual orientation with respect to the true vertical, and tasked them to provide estimates of what they thought was upright. We developed stochastic versions of prominent models of multisensory perception and compared their ability to explain the participants’ responses.

## 2 Methods

### 2.1 Overview

To test the hypothesis outlined above experimentally, we placed 28 participants on a motion platform that was capable of physically rotating them around their naso-occipital axis. While seated on this platform, participants wore a head-mounted display (HMD) fitted with a stereo camera, which was itself mounted on a servo-motor. This setup provided an ‘Alternative-Reality’ (AR) system that could be used to manipulate visual orientation independently from the true gravitational vertical. Participants indicated what they thought was upright for various (in)congruent visual-inertial orientation stimuli. We assessed how well a number of alternative models of spatial orientation could account for participants’ responses by fitting each model to the data and comparing model fit indices.

### 2.2 Ethics Statement

The experimental protocol was approved by the ethical commission of the medical faculty of the Eberhard-Karls University in Tübingen, Germany, reference number 352/2017BO2.

### 2.3 Participants

A total of 31 participants were recruited for the experiment. One participant was not able to complete the experiment due to motion sickness; data from two other participants had to be excluded due to an inability to perform the task. Of the remaining 28 participants, 15 were male. The mean age was 29.9 years, with a standard deviation of 7.6. Participants were compensated for their time at a rate of €8 an hour.

### 2.4 Setup

Visual stimuli were presented using a custom made AR system. This system consisted of an OVRVision Pro stereo camera (Wizapply, Osaka, Japan) mounted via a Dynamixel AX12-A servo motor (Robotis, Lake Forest, California, United States) to a Vive HMD (HTC, New Taipei City, Taiwan). Mounting hardware was designed and 3D-printed in-house. The HMD displayed the images of the left and right camera in the respective screens at a rate of 45 frames per second with a resolution of 1200 × 1080px. The servo allowed us to manipulate the orientation of the participants’ immediate visual surroundings with an accuracy of 0.29°. The software to control the servo and transmit the camera images was also developed in-house. The AR system is shown in Figure 1, left panel.

**Fig. 1:**
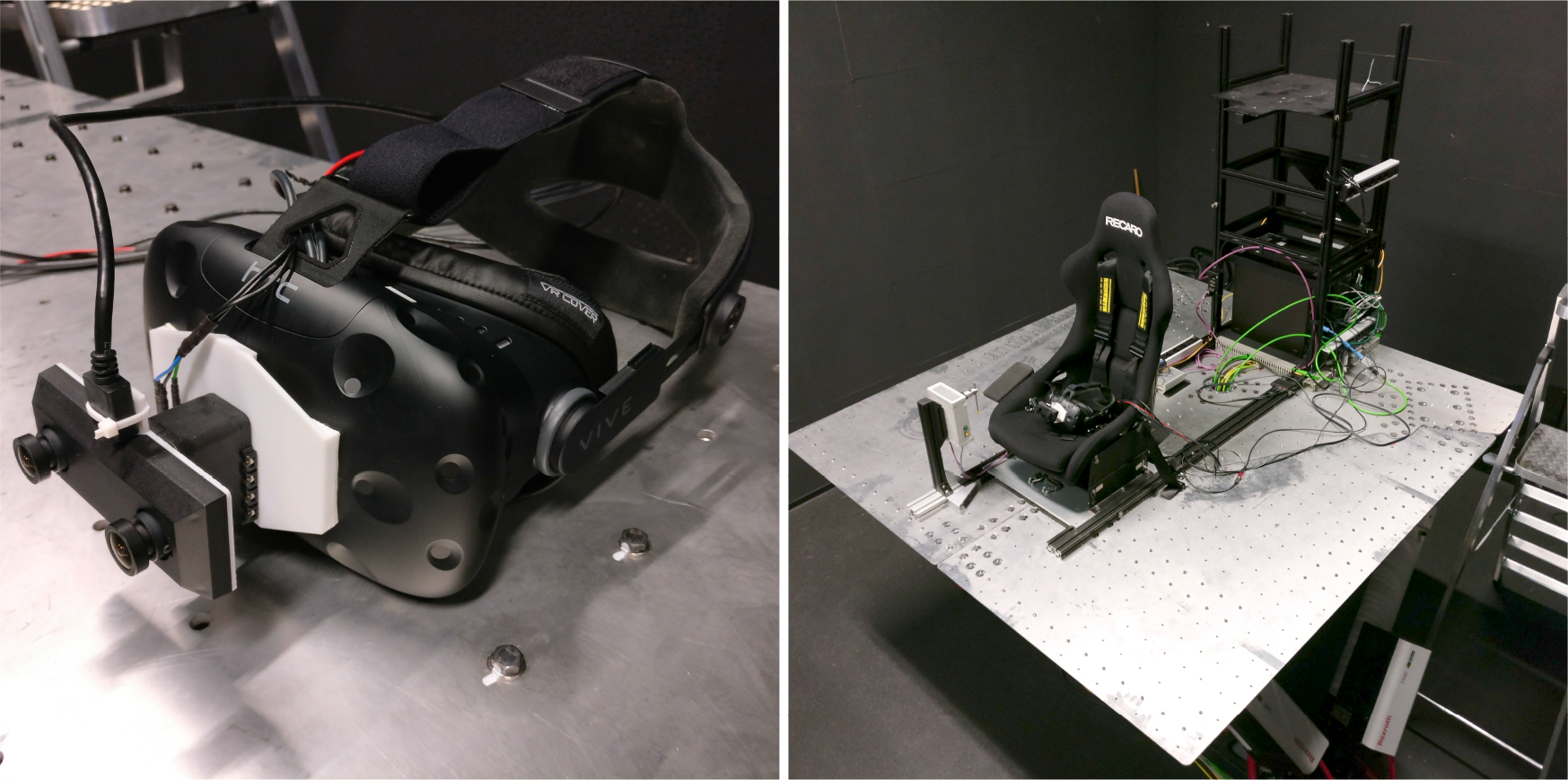
Left panel: photograph of the AR system. Right panel: bird’s eye view of the experimental setup, showing the motion platform and seat. The pointer device is on the right hand side of the seat.

Inertial orientation stimuli were presented using an eMotion 1500 hexapod motion system (Bosch Rexroth AG, Lohr Am Main, Germany). For different physical orientation stimuli, the platform was moved in such a way that the axis of rotation coincided with each participant’s naso-occipital axis. The platform was controlled using Simulink software (The MathWorks, Inc., Natick, Massachusetts, United States).

Participants were seated in an automotive style bucket seat (RECARO GmbH, Stuttgart, Germany) that was mounted on top of the platform, and secured with a 5-point safety harness (SCHROTH Safety Products GmbH, Arnsberg, Germany). To minimize head movements, participants also wore a philadelphia type cervical collar. To ensure that participants could not simply see any tilt of the platform relative to the room, we moved the chair to the edge of the platform. A (monoscopic) screenshot of a participant’s view is presented in supplementary material Figure S1.

To mask the sounds of the motion platform and the servo-motor, participants wore earplugs with a 33dB signal-to-noise ratio (Honeywell Safety Products, Roissy, France) as well as a wireless headset (Plantronics, Santa Cruz, California, United States) that actively canceled outside noise, and that played white noise during platform rotations. A photograph of the complete setup is presented in the right panel of Figure 1.

### 2.5 Task & Stimuli

Participants were tasked to provide estimates of what they believed was the true upright on 400 experimental trials. In each trial, we manipulated the orientation of the visual environment using the AR-system, and the participants’ physical orientation by tilting the motion platform. There were 25 experimental conditions, comprising all possible combinations of visual and physical roll-tilt angles of [–10, –5, 0, 5, 10]°, where 0° corresponds to the gravitational vertical. For participants 20-28, the range of angles was slightly inflated, to [–13, –6.5, 0, 6.5, 13]°. Equal values for the visual and physical roll tilt angle indicate that the camera was aligned with the participants’ physical orientation relative to gravity. Larger visual than physical angles indicate that the camera was tilted more than the platform. For each condition, there were 16 repetitions, totaling 400 trials.

Responses were gathered using a pointer device. This device consisted of a 15cm stainless steel rod mounted to a potentiometer. About 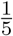 of the length of the rod extended above its center of rotation. This short hand was to be interpreted as the pointer’s top-end, and had to be pointed upwards. The pointer device was free of discontinuities, was not affected by rotation relative to gravity, and provided a < 0.1° resolution. Participants registered a pointer setting as a response by pressing a button at the base of the rod. To ensure the independence of trials, we tested three methods of transitioning from the visual and physical orientations on one trial to the next. For the first nine participants, the camera was turned off while the platform was moved. For the latter five of these participants, heave and sway vibrations were added to the motion profile. These vibrations were in the range of 4 – 8Hz and had a root mean square of approximately 0.1m/s^2^. These vibrations are comparable to road rumble. For the remainder of the participants, the camera was always on during transitions, and there were no vibrations. Regardless of the transition method, the velocity profile of the rotation followed a raised cosine bell, with a duration that was randomly varied between 3 – 4s. Because we did not find any differences in the results of the different subgroups, the data of all participants are presented in a single analysis.

Including instructions and 5-minute breaks every 15 minutes, the experiment lasted between two and three hours.

### 2.6 Models of Spatial Orientation

The aim of the present modeling efforts is to assess the tenability of a number of prominent theories on how the Central Nervous System (CNS) constructs perceptions of upright under conditions of uncertainty about the causality of potential cues on the upright. We identify visual (V) and inertial (I) sensory estimates as likely cues from which a perception of upright may be constructed. The visual system can generate estimates of orientation using polarity information that is present in the optical array (e.g., blue sky/green grass; objects lying on a shelf). Our body’s inertial sensors comprise the vestibular system of the inner ear and various kinds of other sensory neurons distributed throughout the body. Because all these neurons are, either directly or indirectly, responsive to accelerations, we treat them as one single inertial system. The inertial sensory system can generate estimates of orientation by identifying the direction of gravity.

For either sensory modality *m* ∈ {V, I}, we assume that the it generates an estimate *x*_*m*_ of the upright, which is a realization of a random variable that is a possibly distorted version of the respective stimulus orientation *θ*_*m*_. We further assume that for the presently investigated relatively small range of orientations around the true gravitational upright (0 ± 10°), the noise can be approximated by a Gaussian distribution with standard deviation *σ*_*m*_, and that the distortion can be expressed with a scaling parameter *β*_*m*_^1^:

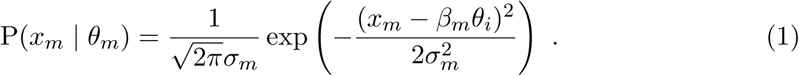

We do not have access to the sensory estimates *x*_*m*_, and are interested in the probability of the responses *r* given stimuli *θ*_*V*_, *θ*_*I*_. This probability is

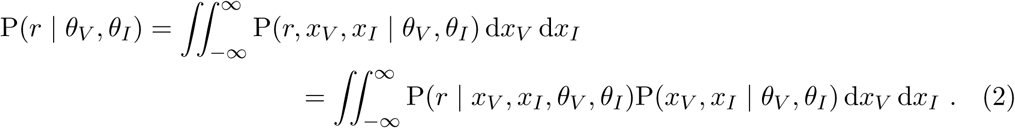

We assume that the sensory estimates for different modalities are generated independently. Therefore, P(*x*_*V*_, *x*_*i*_ | *θ*_*V*_, *θ*_*I*_) = P(*x*_*V*_ | *θ*_*V*_)P(*x*_*I*_ | *θ*_*I*_) and the equation above can be rewritten as

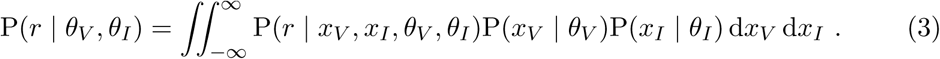

Since *x*_*V*_ and *x*_*I*_ is the only information available to the observer, the response *r* will be conditionally independent of other variables apart from *x*_*V*_ and *x*_*I*_, i.e.,

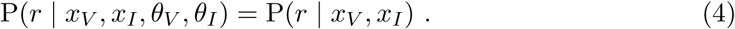

By taking this into account and substituting (1) into (3), we get

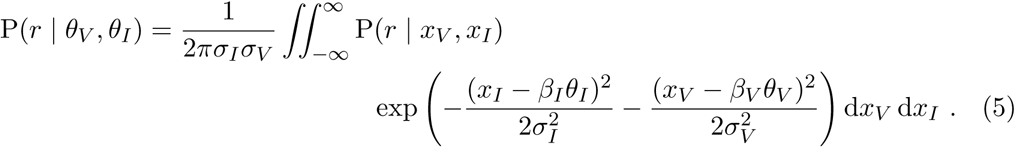

Because a participant is unaware of *θ*_*m*_ and *β*_*m*_, the response generation model P(*r* | *x*_*m*_) uses a different estimate (*x*_*m*_) generation model than the true estimate generation model. From a participant’s perspective, the likelihood of the sensory estimate given any orientation Θ being the true orientation can be expressed as

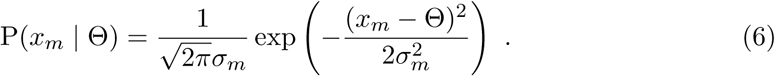

Moreover, we consider the notion that the CNS includes a-priori beliefs about Θ, namely that the upright is usually aligned with the long body axis (above the head), in the construction of this percept. We choose the long body axis as the reference (0°) for other angles. Consequently, we define a prior belief of the following form:

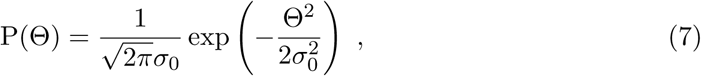

where *σ*_0_ is the distribution’s standard deviation, which represents the strength of the prior belief. We refer to this prior as the ‘idiotropic prior’.

Various general strategies have been proposed on how the CNS may construct perceptions out of the outlined multisensory estimates and prior beliefs. Below, we provide mathematical formulations of prominent strategies, applied to the perception of upright.

#### 2.6.1 Cue Capture

According to Cue Capture (CC) models, perception of specific environmental properties is dominated by a single sensory modality. In the present case, there are two such possibilities: perception of the upright is dominated by either visual or inertial information. A prior belief that the upright aligns with the long body axis interacts with the sensory information according to Bayes’ rule. The posterior probability of Θ given either sensory estimate *x*_*m*_ is then given by:

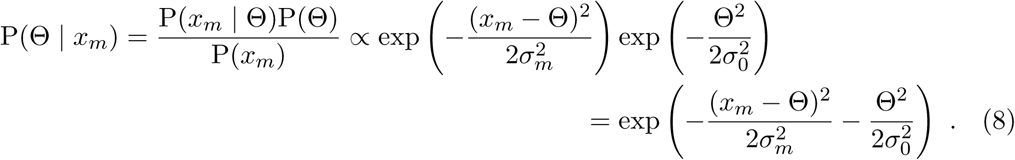

Consistent with the literature, we assume that the response on an individual trial *r* is the mode of this posterior distribution (i.e., the Maximum-A-Posteriori estimate, MAP)

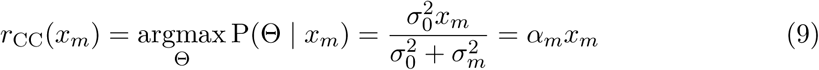

where 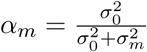. The corresponding PDF can be expressed as

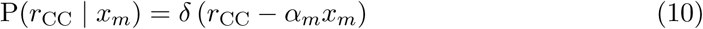

in which *δ*(·) is Dirac’s delta function. By substituting (10) into (5) we obtain

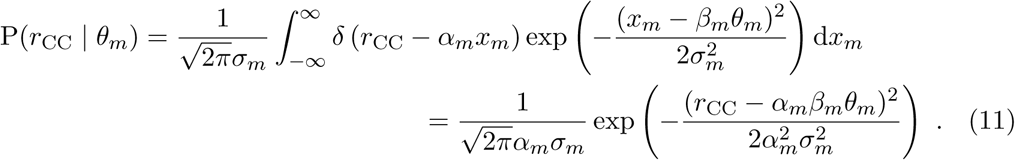

#### Switching Strategy

The CC models can be considered special cases of the Switching Strategy (SS) model. The SS model essentially combines the two alternative CC models: the CNS constructs responses for each sensory modality as in the Cue Capture models, but randomly chooses either modality as dominant source on a trial-by-trial basis:

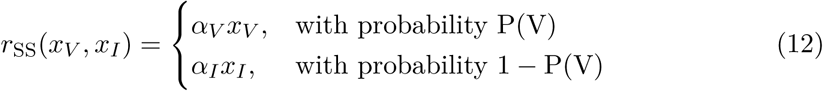

with *α*_*m*_ as before, with the corresponding probability density function

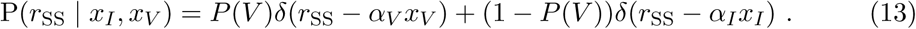

Filling in (13) into (5), we obtain the likelihood of the responses given the stimuli P(rss | *θ*_*V*_, *θ*_*I*_):

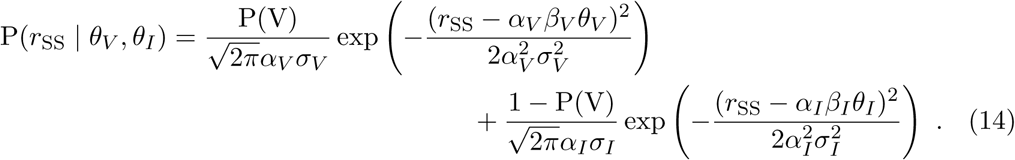

#### 2.6.3 Forced Fusion

In the present formulation of the Forced Fusion (FF) model, it is assumed that visual and inertial sensory estimates are independent from each other, and that both are always interpreted as cues to orientation. The posterior probability of Θ can be expressed as:

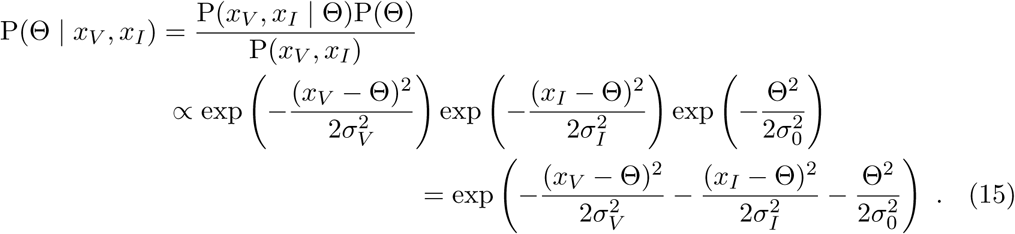

The response *r*_FF_ is again the mode of the posterior distribution (the MAP):

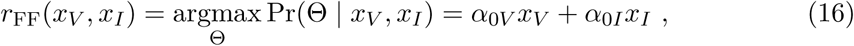

with 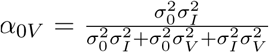, and 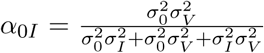. Similar to (10), the posterior PDF can be expressed as

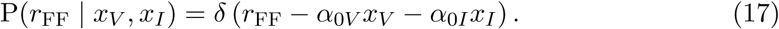

By substituting (17) into (5) and subsequently simplifying, we obtain

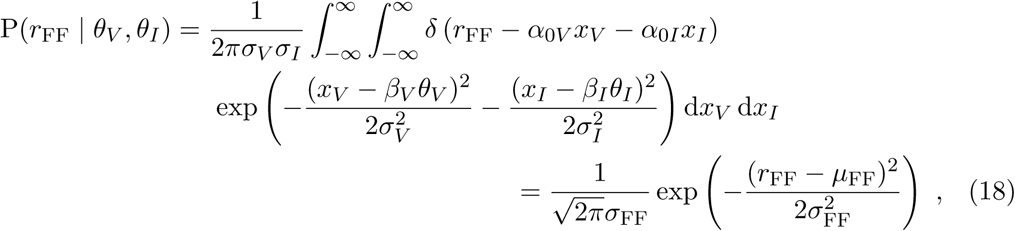

where

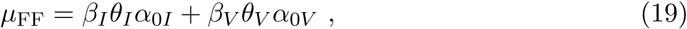

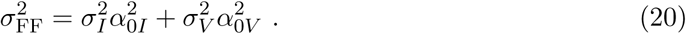

#### 2.6.4 Causal Inference Model

In the models presented above, it is assumed that the CNS either segregates (CC, SS) or fuses multisensory information (FF). CI models allow segregation and fusion to occur in tandem; the estimates generated by the different strategies are treated as intermediate estimates, and a final estimate is constructed by taking into account the probability that the internal estimates share a common cause (C), favoring FF; or have independent causes (C̅), favoring SS.

The probability of a common cause given the sensory estimates is

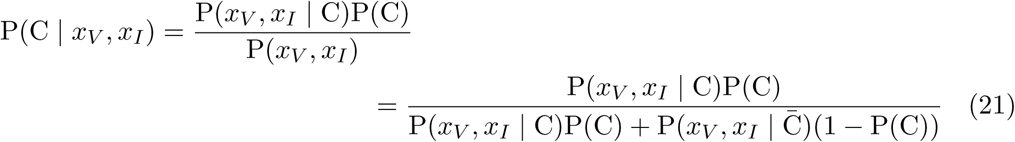

where P(C) is a free parameter that represents a prior tolerance for discrepancies. The likelihood of the sensory estimates *x*_*V*_, *x*_*I*_ given a common cause C is

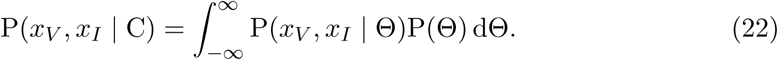

where P(Θ) is the idiotropic prior. P(*x*_*V*_, *x*_*I*_ | Θ) is the likelihood of *x*_*v*_, *x*_*I*_ given some common orientation Θ. This becomes

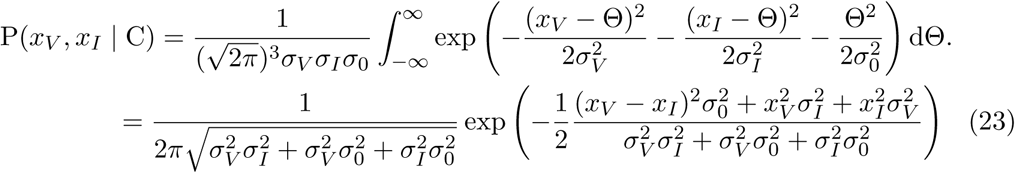

For an interpretation of independent causes, a response will be based on either the visual or the inertial estimate. We assume that the same idiotropic prior interacts with both sensory estimates, as only one of them will ultimately be treated as informative of the upright.

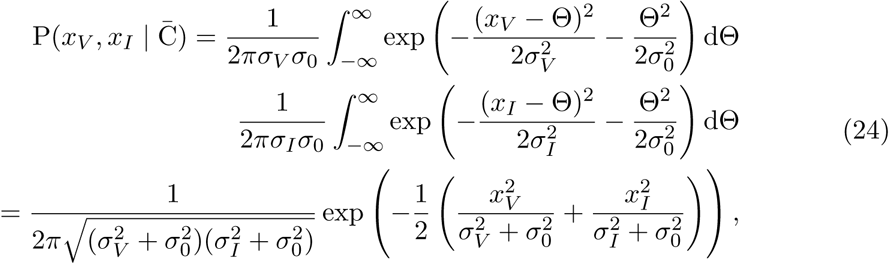

As in [18, 19], the response *r*_CI_ is a weighted average of the response according to the SS strategy *r*_ss_ and the response according to the FF strategy *r*_FF_, with weights proportional to the respective probability of the causal structures

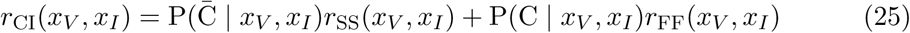

where P(C̅ | *x*_*V*_, *x*_*I*_)) = 1 – P(C | *x*_*V*_, *x*_*I*_). *r*_FF_ is a deterministic function of (*x*_*V*_, *x*_*I*_) and *r*_SS_ is a random variable which can take one of the two values according to (12). Therefore, rei can be written as

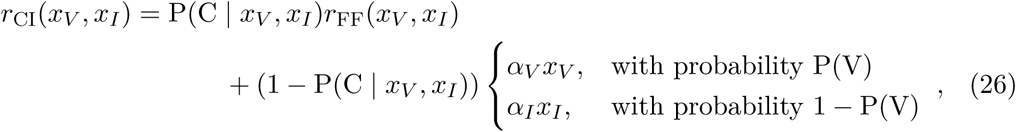

which can be expressed as the density function

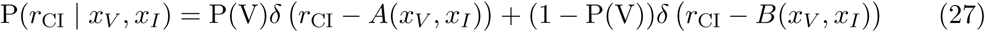

where

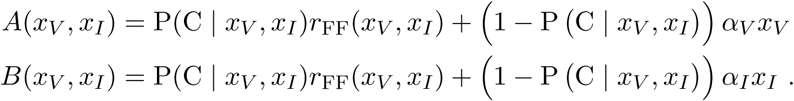

By substituting the expression for *r*_FF_ (16) into the equations above and doing some transformations we obtain

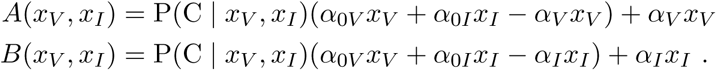

By replacing the response generating model P(*r* | *x*_*V*_, *x*_*I*_) in (5) with (27), we can obtain the likelihood function for the responses given the stimuli. However, due to the P(C | *x*_*V*_, *x*_*I*_) expression, the integral in (5) cannot be represented in a closed form. To resolve this issue, we linearize *A*(*x*_*V*_, *x*_*I*_) and *B*(*x*_*V*_, *x*_*I*_) at *x*_*V*_ = *x*_*V*0_ = *β*_*V*_*θ*_*V*_, *x*_*I*_ = *x*_*I*0_ = *β*_*I*_ *θ*_*I*_. For *A*, *B*, we obtain

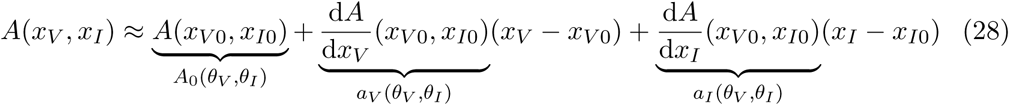

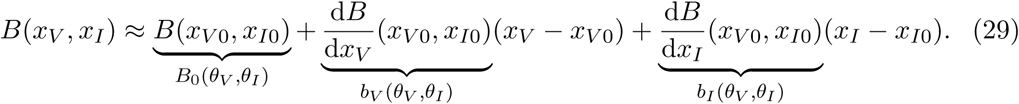

If we use the approximations for *A*(*x*_*V*_, *x*_*I*_), *B*(*x*_*V*_, *x*_*I*_) in (27), we can solve the integrals in (5). This yields

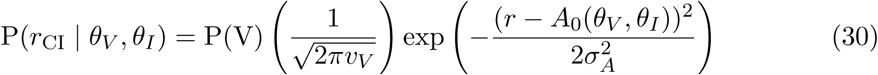

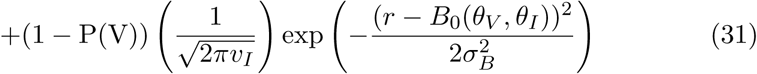

with 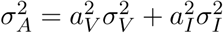 and 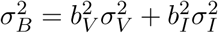.

We tested whether the linear approximation was appropriate by comparing the results with those of performing numerical integrations for the initial few participants.

### 2.7 Data analysis

For each of the models described in the Models of Spatial Orientation section above, the fit of two versions was assessed.

In the first version of the models, scaling of perceptions was modeled with parameters *β*_*V*_ and *β*_*I*_ (as described above). In the second version of the models, the values of the *β*_*V*_ and *β*_*I*_ parameters were fixed at 1, reflecting an assumption that perception itself is veridical. Instead, here distortion of the final response *R* was attributed to an over-or underestimation of the angle of the rod. For the CC and FF models, this was implemented as a linear transformation of the response random variable: *R* = *β*_*r*_*r*_model_, with 𝔼(*R*) = 𝔼(*β*_*r*_*r*_model_) = *β*_*r*_𝔼(*r*_model_), and 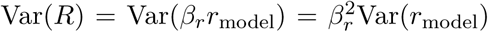. For the SS and CI models, scaling by *β*_*r*_ was applied to the mixture components individually.

The parameters to account for distortions (*β*_*V*_, *β*_*I*_, or *β*_*r*_); the standard deviations of the sensory estimates (*σ*_*V*_, *σ*_*I*_); and the mixture and ‘tolerance for discrepancies’ prior parameters (P(V), P(C)) were treated as free parameters. The standard deviation of the idiotropic prior σο was fixed a value of 100. The value needed to be fixed to ensure that the optimization procedures could properly converge, and the large value of 100 was chosen to reflect the observation that the prior has a negligible effect on the subjective vertical[20, 17]. This renders the prior effectively uninformative, but we chose to include it for consistency with the literature.

It should be noted that while it is theoretically possible that distortions are introduced both in the perceptual process as well as in the translation from percepts to responses, it was not possible to estimate both effects simultaneously. This is due to the fact that perceptual and response distortion parameters allow the models to produce similar behavior, thus resulting in an infinite number of equivalent solutions when both possibilities are allowed.

We fitted the CC and FF models by minimizing the negative log-likelihood of the responses given the model 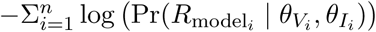;, using the fmincon routine in MATLAB. The fmincon routine was not suitable to fit the SS and CI models: these are mixture models, and directly maximizing the likelihood can lead to numerical issues. We therefore applied the Expectation-Maximization (EM) algorithm to fit these models [21]. In the EM-algorithm, membership of mixture components is treated as a latent variable. The model likelihood is maximized iteratively, by repeating a set of two steps: first, the probability of each observation belonging to either component of the mixture is determined given an initial set of parameters. Second, the model parameters are re-estimated, while taking the probability of component membership calculated in the previous step into account. To re-estimate the parameters in the second step, we again minimized the model negative log-likelihoods using the MATLAB fmincon routine. The iterations were terminated when the change in model likelihood was smaller than 1E-6. To determine which model best approximated participant responses, we compared model Bayesian Information Criterion (BIC) scores [22]. The BIC is a penalized likelihood score, taking into account the number of observations and the number of free parameters in each model. The model with the lowest BIC score is considered the best in an absolute sense. Differences in model BIC scores (ΔBIC) between 0-2; 2-6; 6-10 are considered negligible, positive, and strong evidence, respectively, and ΔBIC > 10 are considered decisive evidence.

## 3 Results

A comparison of the fit of the two different versions of the models (where either perceptions or responses were allowed to exhibit distortions, as described in the Data analysis section) showed that the results were almost identical. Because the models with a single scaling parameter for the response are simpler than dual scaling parameters for the perceptions, we conclude that visual and inertial perceptions of orientation are veridical. Consequently, we report here only the results of the models with a scaling of the response. All optimization results are available as supplementary material: results for the first version of the models (scaling of perception) are provided as supplementary material Table S1; the results of the second versions (scaling of responses) are provided in supplementary material Table S2.

### 3.1 Model comparisons

The model fitting procedure was performed for each participant individually. As an illustration, Figure 2 shows the data and model fits for an example participant.

**Fig. 2:**
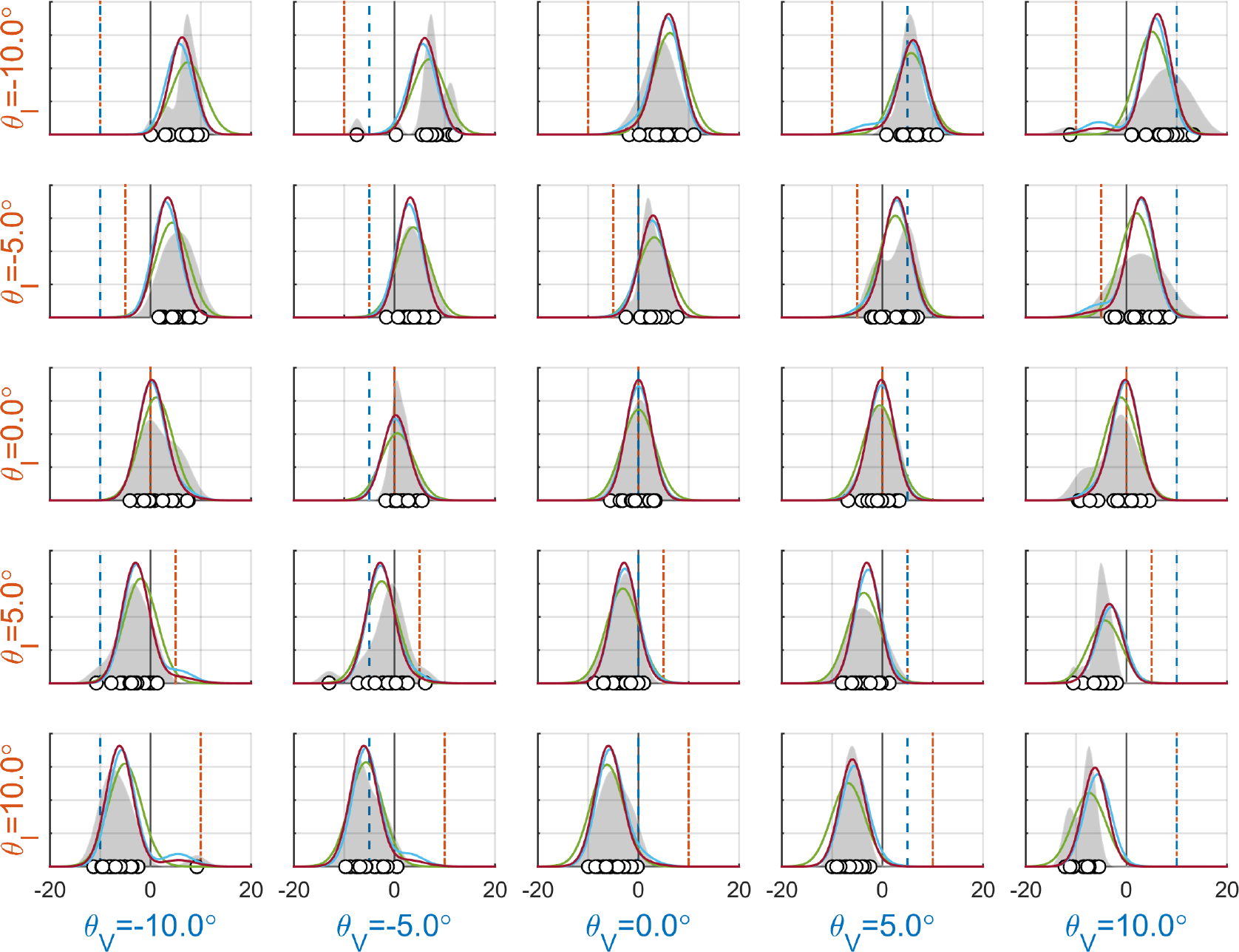
Overview of the results for an exemplary participant (8). Each panel shows the results for a particular experimental condition. The thin black line at 0° is the Earthvertical. The dashed lines show the visual (light-blue) and physical (orange) tilt angle. The white dots show individual responses; the gray-shaded areas show the corresponding kernel density estimates. The colored lines represent the densities according to the SS (blue), FF (green), and CI (red) models. Note that 1. participants were instructed to point upwards, and that responses therefore reflect the inverse of the perceived tilt; 2. the kernel density estimates in each panel are based on 16 data points.

Combined over all participants, the evidence favors the FF model, as indicated by a ΔBIC of 102.90 compared to the runner-up model, CI. However, a closer inspection of the individual results provides considerable evidence that the preferred model differs between individuals. Specifically, compared to the runner-up models, a variant of the cue capture model (either CC_I_ or SS) explained the data best for 9 participants, with evidence mostly positive (median ΔBIC = 3.96); the FF model explained the data best for 12 participants, with mostly strong evidence (median ΔBIC = 6.28); and the CI model provides the best explanation for 7 participants, with mostly decisive evidence (median ΔBIC = 13.80). Model BIC scores for all participants are presented in Table 1.

**Table 1:** BIC scores for models of estimation of the vertical. The best BIC score is boldfaced for each participant (pp). For each model, overall BIC scores were calculated using the sums (over participants) of the model log-likelihoods, the number of parameters, and the total number of observations.

| pp | BIC |  |  |  |  |
| --- | --- | --- | --- | --- | --- |
|  | CC<sub>V</sub> | CC<sub>I</sub> | SS | FF | CI |
| 1 | 3077.53 | **2166.61** | 2173.01 | 2168.54 | 2174.69 |
| 2 | 3076.08 | 2359.88 | 2368.18 | **2354.20** | 2362.49 |
| 3 | 3104.46 | 2787.09 | **2753.21** | 2790.37 | 2757.37 |
| 4 | 2897.09 | 2218.88 | 2227.18 | **2213.62** | 2221.90 |
| 5 | 3705.19 | **2980.61** | 2988.92 | 2983.59 | 2991.90 |
| 6 | 2889.75 | 2585.14 | 2593.45 | 2502.99 | **2500.83** |
| 7 | 3236.36 | **2670.09** | 2678.39 | 2674.24 | 2678.58 |
| 8 | 2594.37 | 2120.68 | 2076.28 | 2101.50 | **2053.25** |
| 9 | 2666.04 | 2153.48 | 2161.79 | **2080.68** | 2087.57 |
| 10 | 3092.81 | **2174.52** | 2182.83 | 2175.84 | 2184.13 |
| 11 | 2540.78 | 2446.51 | 2280.46 | 2310.40 | **2242.21** |
| 12 | 2754.36 | 2321.96 | **2320.33** | 2321.32 | 2320.80 |
| 13 | 2647.50 | 2127.72 | 2136.02 | **2077.34** | 2084.60 |
| 14 | 2722.95 | 2084.13 | 2092.44 | **2036.48** | 2044.79 |
| 15 | 2882.74 | **2466.69** | 2475.00 | 2470.85 | 2473.72 |
| 16 | 2714.79 | 2682.75 | 2610.27 | **2567.65** | 2570.66 |
| 17 | 2553.42 | 2076.06 | 2053.34 | 2015.75 | **2000.14** |
| 18 | 2775.11 | 2443.46 | 2433.93 | **2368.28** | 2376.37 |
| 19 | 2689.46 | 2210.52 | 2218.82 | 2194.40 | **2194.22** |
| 20 | 3161.85 | 2503.30 | **2494.27** | 2507.46 | 2498.23 |
| 21 | 2893.19 | 2511.96 | 2520.27 | **2493.97** | 2502.09 |
| 22 | 3133.62 | 3093.23 | 3070.04 | **3066.07** | 3073.49 |
| 23 | 3333.31 | 2752.27 | 2760.58 | 2749.66 | **2745.67** |
| 24 | 2571.22 | 2173.98 | 2182.29 | **2084.22** | 2091.47 |
| 25 | 2893.14 | 2304.81 | 2310.93 | **2290.21** | 2292.17 |
| 26 | 2823.62 | 2546.91 | 2508.35 | 2495.64 | **2481.84** |
| 27 | 3213.84 | **2914.54** | 2919.08 | 2918.68 | 2923.21 |
| 28 | 2489.34 | 2690.98 | 2463.29 | **2450.49** | 2452.35 |
| overall | 81320.63 | 68755.51 | 68635.82 | **67953.97** | 68056.87 |

### 3.2 Parameter estimates

Given the between-subject variability in which model best explained the data, we consider the obtained parameter estimates separately for each model. A summary of the parameter estimates is provided in Table 2.

**Table 2:** Median values (and standard deviation) of parameter estimates for each model. Except for the CC_V_ model, the values presented here are calculated using only the parameters obtained for those participants for whom the respective model provided the best explanation of the data (as per Table 1). The parameters for the CC_v_ model are median values over all participants instead, as this model never provided the best fit. The lower bounds of 1 for **σ**_*m*_ were chosen to prevent cases where fitting of the mixture models would result in explanation of a single outlier with a dedicated component with near-zero standard deviation.

| parameter | model |  |  |  |  | bounds |
| --- | --- | --- | --- | --- | --- | --- |
| | $CC_V$ | $CC_I$ | SS | FF | CI | |
| $\beta_r$ | 0.92(0.38) | 1.54(0.82) | 1.13(0.27) | 1.15(0.39) | 0.86(0.30) | [0.5-5] |
| $\sigma_V$ | 20.18(13.98) | | 12.57(6.94) | 10.80(4.06) | 8.09(3.74) | [1-∞] |
| $\sigma_I$ | | 3.44(2.26) | 5.15(0.42) | 4.88(6.01) | 4.36(0.87) | [1-∞] |
| P(V) |  |  | 0.02(0.06) |  | 0.23(0.38) | [0-1] |
| P(C) |  |  |  |  | 0.28(0.19) | [0-1] |

The scaling of responses, as expressed by parameter *β*_*r*_, was generally close to unity, although for some participants values considerably larger or smaller than one were estimated. A value larger than one indicates that responses showed overestimation, or equivalently, that a participant underestimated the tilt angle of the rod.

Median values for parameters *σ*_*V*_ and *σ*_*I*_ ranged between 8.09 – 20.18°, and 3.44 – 5.15°, respectively. This indicates that visual estimates of the upright were roughly twice as uncertain as inertial estimates, or, assuming fusion of sensory estimates, that the contribution of inertial information to the construction of perceptions of the upright is roughly four times as big as the contribution of visual information.

For the SS and CI models, parameter P(V) was found to be close, but not equal, to zero for the SS model (median 0.02), suggesting that these participants generally relied on inertial information (as in the CC_I_ model), but occasionally lapsed by reporting the visual tilt angle. In the CI model, the value of this parameter was larger (median 0.23), showing a larger probability that responses reflected visual information when a discrepancy was detected.

The a-priori tolerance for discrepancies parameter P(C) of the CI model had a median value of 0.28, suggesting a stronger tendency to attribute cues to independent causes than to assume a common cause.

A complete overview of the obtained parameter estimates is available in supplementary materials Table S1 and S2.

## 4 Discussion

The present study was designed to assess how the subjective experience of the upright is constructed from visual and inertial information, with a particular focus on how this process is affected by discrepancies between the different cues to the upright.

We placed participants on a motion platform, and presented them with various (discrepant) combinations of visual and physical tilt angles. To manipulate the visual tilt independently from the physical tilt while maintaining the highest degree of ecological validity possible, we developed an Alternative Reality (AR) system. The AR system consists of a stereo camera mounted via a servo motor to an HMD. Using this setup, we were able to manipulate the tilt of our participants’ actual visual surroundings. Importantly, this implies that it was impossible for participants to know whether the visual cue, at any one moment, provided veridical information or not[16].

For each combination of tilt angles, we asked our participants to indicate the upright. We then compared how well a number of prominent models of multisensory perception from the literature could explain the findings. The models that were considered can be regarded as forming a spectrum of possible strategies the central nervous system (CNS) could apply to form estimates of any particular environmental property out of multiple potential sources of information on this property. One end of the spectrum is formed by Cue Capture (CC) and Switching Strategy (SS) models. In CC models[23], it is assumed that multisensory perception is dominated by a single sensory modality; similarly, SS models assume that perception is dominated by a single modality, but the dominant modality is allowed to alternate on a trial-by-trial basis[24, 25]. The other end of the spectrum is formed by Forced Fusion (FF) models[9, 10]. Here, it is assumed that all modalities involved provide information on the property of interest, and a multisensory estimate is formed by determining the most likely single value for the property, given the sensory estimates and prior beliefs about the property. In between these extremes come Causal Inference (CI) models [18]. According to CI, the CNS includes inferences on the causality of multisensory information in the process of forming estimates of a property: essentially treating the estimates according to FF and CC/SS as intermediates, and choosing between them or weighting them according to the inference made on the signals’ causality[26].

We developed statistical formulations of each of these models, and compared them on the basis of the quality of their fits to participants’ responses in a task where they indicated the ‘true’ (physical) upright, while being exposed to incongruent visual and physical tilt. Overall, the results decide in favor of the FF model. This implies that the CNS automatically fuses multisensory information on spatial orientation, at least for the range of discrepancies tested (up to ±27°). This is surprising, considering that these discrepancies are much larger than the size of the noises of visual and inertial estimates (*σ*_*V*_ ≈ 8°; *σ*_*I*_ ≈ 4°).

Although the conclusion that perception of the upright is described by FF would be consistent with several earlier studies[11, 11, 14, 17, 13], a closer look at the individual data reveals that there is a substantial variability with respect to which model best explains the data. Moreover, the evidence to discriminate between models ranges between positive and decisive in the majority of the cases (22/28), which justifies further consideration of these differences. Specifically, we find that for 9/28 participants, the CC/SS model explains the data best; for 12/28 participants the FF model fits best; and for the remaining 7/28 the CI model provides the best explanation.

We contend that the results of the present study resemble those of recent work on CI in the heading estimation. According to Einstein’s equivalence principle [27], gravity is equivalent to a constant acceleration. Our orientation relative to gravity can therefore be considered analogous to the heading of horizontal linear accelerations. Given that this premise is accepted, the present findings are similar to those of two recent studies on estimation of heading of translational motions in the horizontal plane [28, 29]. Findings of the earlier study favored FF ‒although evidence for CI was found for one participant[28]. In the subsequent study[29], the authors adopted a more elaborate experimental paradigm, and obtained decisive evidence in favor of CI. Because the findings of the present study populate the spectrum of models more or less evenly, it is possible that the chosen range of discrepancies was too narrow to reveal segregation behavior for some participants, whereas it may have been too broad for others, thereby concealing fusion behavior. Consequently, it is likely that future work, using a more refined experimental paradigm, will yield more definitive proof for a form of CI.

## Footnotes

1 we corrected for bias (constant offset) in a participant’s responses at the outset of the experiment. This was done by calibrating the alignment of the rod such that participants’ subjective upright in the absence of any experimental manipulations yielded a pointer device output of zero, and also by subtracting from each data point the average response.

## References

[1] I. P. Howard, S. S. Bergstrom, M. Ohmi, Shape from shading in different frames of reference, Perception 19 (4) (1990) 523–530.

[2] I. P. Howard, Human visual orientation, John Wiley & Sons, 1982.

[3] J. R. Lackner, A. Graybiel, Postural illusions experienced during z-axis recumbent rotation and their dependence upon somatosensory stimulation of the body surface., Aviation, space, and environmental medicine.

[4] J. R. Lackner, A. Graybiel, Some influences of touch and pressure cues on human spatial orientation., Aviation, space, and environmental medicine.

[5] D. Vaitl, H. Mittelstaedt, F. Baisch, Shifts in blood volume alter the perception of posture, International Journal of Psychophysiology 27 (2) (1997) 99–105.

[6] D. Vaitl, H. Mittelstaedt, R. Saborowski, R. Stark, F. Baisch, Shifts in blood volume alter the perception of posture: further evidence for somatic graviception, International Journal of Psychophysiology 44 (1) (2002) 1–11.

[7] H. Mittelstaedt, Somatic graviception, Biological psychology 42 (1) (1996) 53–74.

[8] H. Mittelstaedt, A new solution to the problem of the subjective vertical, Naturwissenschaften 70 (6) (1983) 272–281.

[9] M. O. Ernst, M. S. Banks, Humans integrate visual and haptic information in a statistically optimal fashion, Nature 415 (6870) (2002) 429–433.

[10] M. O. Ernst, H. H. Hülthoff, Merging the senses into a robust percept, Trends in cognitive sciences 8 (4) (2004) 162–169.

[11] R. T. Dyde, M. R. Jenkin, L. R. Harris, The subjective visual vertical and the perceptual upright, Experimental Brain Research 173 (4) (2006) 612–622.

[12] R. A. A. Vingerhoets, M. De Vrijer, J. A. A Gisbergen, W. P. Medendorp, Fusion of visual and vestibular tilt cues in the perception of visual vertical, Journal of neurophysiology 101 (3) (2009) 1321–1333.

[13] I. A. Clemens, M. De Vrijer, L. P. Selen, J. A. A Gisbergen, W. P. Medendorp, Multisensory processing in spatial orientation: an inverse probabilistic approach, Journal of Neuroscience 31 (14) (2011) 5365–5377.

[14] L. R. Harris, M. Jenkin, H. Jenkin, J. E. Zacher, R. T. Dyde, The effect of longterm exposure to microgravity on the perception of upright, npj Microgravity 3 (1) (2017) 3.

[15] R. T. Dyde, M. R. Jenkin, H. L. Jenkin, J. E. Zacher, L. R. Harris, The effect of altered gravity states on the perception of orientation, Experimental brain research 194 (4) (2009) 647–660.

[16] K. N. de Winkel, G. Clement, E. L. Groen, P. J. Werkhoven, The perception of verticality in lunar and martian gravity conditions, Neuroscience letters 529 (1) (2012) 7–11.

[17] R. A. A. Vingerhoets, W. P. Medendorp, J. A. A Gisbergen, Body-tilt and visual verticality perception during multiple cycles of roll rotation, Journal of neurophysiology 99 (5) (2008) 2264–2280.

[18] K. P. Körding, U. Beierholm, W. J. Ma, S. Quartz, J. B. Tenenbaum, L. Shams, Causal inference in multisensory perception, PLoS ONE 2 (9) (2007) e943.

[19] U. Beierholm, L. Shams, W. J. Ma, K. Koerding, Comparing bayesian models for multisensory cue combination without mandatory integration, in: Advances in neural information processing systems, 2008, pp. 81–88.

[20] S. B. Bortolami, A. Pierobon, P. DiZio, J. R. Lackner, Localization of the subjective vertical during roll, pitch, and recumbent yaw body tilt, Experimental brain research 173 (3) (2006) 364–373.

[21] A. P. Dempster, N. M. Laird, D. B. Rubin, Maximum likelihood from incomplete data via the em algorithm, Journal of the royal statistical society. Series B (methodological) (1977) 1–38.

[22] G. Schwarz, et al., Estimating the dimension of a model, The annals of statistics 6 (2) (1978) 461–464.

[23] I. Rock, J. Victor, Vision and touch: An experimentally created conflict between the two senses, Science 143 (3606) (1964) 594–596.

[24] K. De Winkel, F. Soyka, M. Barnett-Cowan, H. Bülthoff, E. Groen, P. Werkhoven, Integration of visual and inertial cues in the perception of angular self-motion, Experimental brain research 231 (2) (2013) 209–218.

[25] C. M. Goeke, S. Planera, H. Finger, P. König, Bayesian alternation during tactile augmentation, Frontiers in behavioral neuroscience 10.

[26] D. R. Wozny, U. R. Beierholm, L. Shams, Probability matching as a computational strategy used in perception, PLoS Computational Biology 6 (8) (2010) e1000871.

[27] A. Einstein, Über das relativitätsprinzip und die aus demselben gezogenen folgerungen, Jahrbuch der Radioaktivität und Elektronik 4 (1908) 411–462.

[28] K. N. De Winkel, M. Katliar, H. H. Bülthoff, Forced fusion in multisensory heading estimation, PLoS One 10 (5) (2015) e0127104.

[29] K. N. De Winkel, M. Katliar, H. H. Bülthoff, Causal inference in multisensory heading estimation, PloS one 12 (1) (2017) e0169676.

